# Dengue virus clinical isolates sustain viability of infected hepatic cells by counteracting apoptosis-mediated DNA breakage

**DOI:** 10.1101/2020.06.19.162479

**Authors:** Himadri Nath, Anisa Ghosh, Keya Basu, Abhishek De, Subhajit Biswas

**Affiliations:** Infectious Diseases & Immunology Division, CSIR-Indian Institute of Chemical Biology (IICB), 4, Raja S.C. Mallick Rd, Jadavpur, Kolkata-700032, West Bengal, India; Department of Pathology, Institute of Post-Graduate Medical Education and Research (IPGMER) and Seth Sukhlal Karnani Memorial (SSKM) Hospital, 244, Acharya Jagadish Chandra Bose Rd, Bhowanipore, Kolkata-700020, West Bengal, India; Department of Dermatology, Calcutta National Medical College & Hospital, Kolkata-700014, West Bengal, India; Academy of Scientific and Innovative Research (AcSIR), Ghaziabad, India

**Keywords:** apoptosis, dengue virus, cell free DNA, NS1, virotoxin

## Abstract

NS1, a virotoxin, abundantly present in Dengue patients’ blood, is a major player behind disease patho-biogenesis including plasma leakage and damage to the liver. Despite the presence of NS1 in blood, Dengue is asymptomatic and self-limiting in ≥80% Dengue virus (DV) infected people. We investigated this observation and found that plasmid-mediated NS1 expression and secretion in liver cells (Huh7) are sufficient to cause programmed cell death (apoptosis) and associated cellular DNA breakage. However, liver or kidney cell lines infected with DV and secreting equivalent amounts of NS1 didn’t exhibit apoptotic DNA breakage. In fact, DV-infected cells showed better survival than cells in which only NS1 was transiently expressed by transfection. We also found that DV can even prevent chemical-induced apoptotic DNA damage in infected host cells. So, DV thwarts host antiviral defence i.e. apoptosis, by counteracting cellular DNA breakages and keeps the infected cells metabolically active to prolong virus replication.

## Introduction

Dengue is the most serious life-threatening vector-borne viral disease plaguing mankind. It affects approximately 390 million (95% Credible Interval (CI): 284-528) people annually across the globe of which 96 million (CI: 67-136) manifests clinically. So, there is a large population where Dengue virus (DV) infection is asymptomatic. Dengue is endemic to more than 100 countries which makes one-third of the world’s population to live in risk areas, which are tropical and subtropical regions (1). Dengue virus is a member of the family *Flaviviridae*, genus Flavivirus, and is transmitted to humans by the bite of female *Aedes* mosquitoes. The infection with any of the main four DV serotypes (1–4) can either be asymptomatic or manifest in three clinical forms of increasing severity: Dengue fever (DF), Dengue haemorrhagic fever (DHF) and Dengue shock syndrome (DSS). Dengue fever is characterized by fever, headache, muscle and joint pain and rash. Dengue haemorrhagic fever (DHF) is characterized by thrombocytopenia, increase in vascular permeability and plasma leakage. In severe cases, circulatory failure and shock (DSS) and even death may occur (2). Dengue virus particle is approximately 500Å in diameter. It has a positive sense RNA genome of ~10.7 Kb coding for three structural and seven non-structural proteins. Capsid, precursor-Membrane and Envelope are three structural proteins. Non-structural proteins are NS1, NS2a, NS2b, NS3, NS4a, NS4b and NS5 (3).

Dengue virus infections affect multiple organ systems, the commonest being the liver (4). Hepatic dysfunction is an important feature of Dengue as is evident from various clinical reports (5). Liver involvement in Dengue cases may vary from asymptomatic elevation of hepatic transaminases to severe manifestations in the form of acute liver failure (6). Autopsy of Dengue patients in Myanmar showed damage to liver with moderate to severe sinusoidal congestions involving midzonal and centrilobular areas (7). In a study comprising 240 patients, hepatic dysfunctions in the form of deranged total bilirubin (19.5%), AST (97.7%), ALT (93.9%), ALP (32.6%), albumin (29.1%) and PTI (15.5%) were observed (8).

DV NS1 (48 KDa) is secreted from infected mammalian cells as hexamer and widely used as a diagnostic marker (9). NS1 is essential for virus replication (10,11) and the secreted form contributes to complement fixation (12) and pathogenesis. NS1 has been shown to act as PAMP and activate TLR4, leading to induction and release of pro-inflammatory cytokines and chemokines (13). Circulating levels of soluble NS1 (sNS1) vary from 0.04-0.6 μg/mL for DF, 0.6-2.5 μg/ml with reports of up to 15 μg/mL in case of DHF and persist for up to 4-6 days from the onset of fever (14). These data are based on NS1 levels measured in the bloodstream of infected patients. In mouse model, sNS1 was found to accumulate in liver and the hepatocytes appeared to be the major target cells *in vivo* (15). Internalization and stability of sNS1in human hepatocyte cell lines like HepG2 and Huh7 had also been reported (15). Antibodies against NS1 have been found to promote apoptosis of liver cells in mouse model (16). But the direct effect of clinical DV infections on liver cells in terms of apoptosis was not clear. So, we have studied apoptosis in Huh7 cells upon infection with DV clinical isolates of serotype 1, 2 and 3 not passaged more than four times in cell culture. NS1 secretion in case of these clinical isolates in hepatocyte cells, has been studied in details. African green monkey kidney cell line (Vero) is also used in this study.

Here we estimated sNS1 in cell lines (Huh7, Vero) infected with known copy numbers of DV clinical isolates. Based on the sNS1 levels, NS1 plasmid constructs were transfected in order to express comparable amounts of sNS1in same cell lines to observe the effect of equivalent amounts of NS1 on cells, in presence and absence of DV infection for all the three serotypes.

It was observed that DV infection or NS1 expression, either condition, induces apoptosis in both cell lines. But DV infection exerts control/check over late apoptotic DNA breaks which kept the cells alive and capable of more virus replication. Thus, there exists DV-mediated protection of host cellular DNA from apoptosis-induced breakage. We believe that this is a strategy, DV deploys to delay cell death, thus allowing more time and space for virus replication within the infected cells. Our observation also provides a plausible explanation for the high cell-free DNA that has been routinely observed clinically in DV infections (17).

## Results

### Dengue virus and NS1 only, both can induce Cleaved Caspase3 (CC3)

Secreted NS1 from DV1-HNSB-P4-infected (10 virus genome Equivalents (gE) /cell) and NS1-plasmid-transfected Huh7 cells in six well plates (approx. 10^6^ cells at confluence), were measured in multiple experiments (Supplementary Table1). It was estimated that 1.0 μg pcDNA3.1-NS1 expression construct results in expression of 2.55 (± 0.59) μg/ml DV1-NS1 in the cell supernatant at 96h post-transfection. This was equivalent to 2.53 (± 0.79) μg/ml, expressed at 96h post-infection when the same number of cells were infected with the aforesaid virus inoculum (10 gE/cell). One more NS1-plasmid quantity i.e. 3.3 μg was tested, expressing 4.36 (± 0.33) μg/ml NS1 under similar conditions. The expression levels of CC3 rose with increasing levels of NS1 secretion, as observed in case of 1.0 μg and 3.3 μg NS1 plasmid transfected cells (Fig. 1a, b).

**Fig 1.**
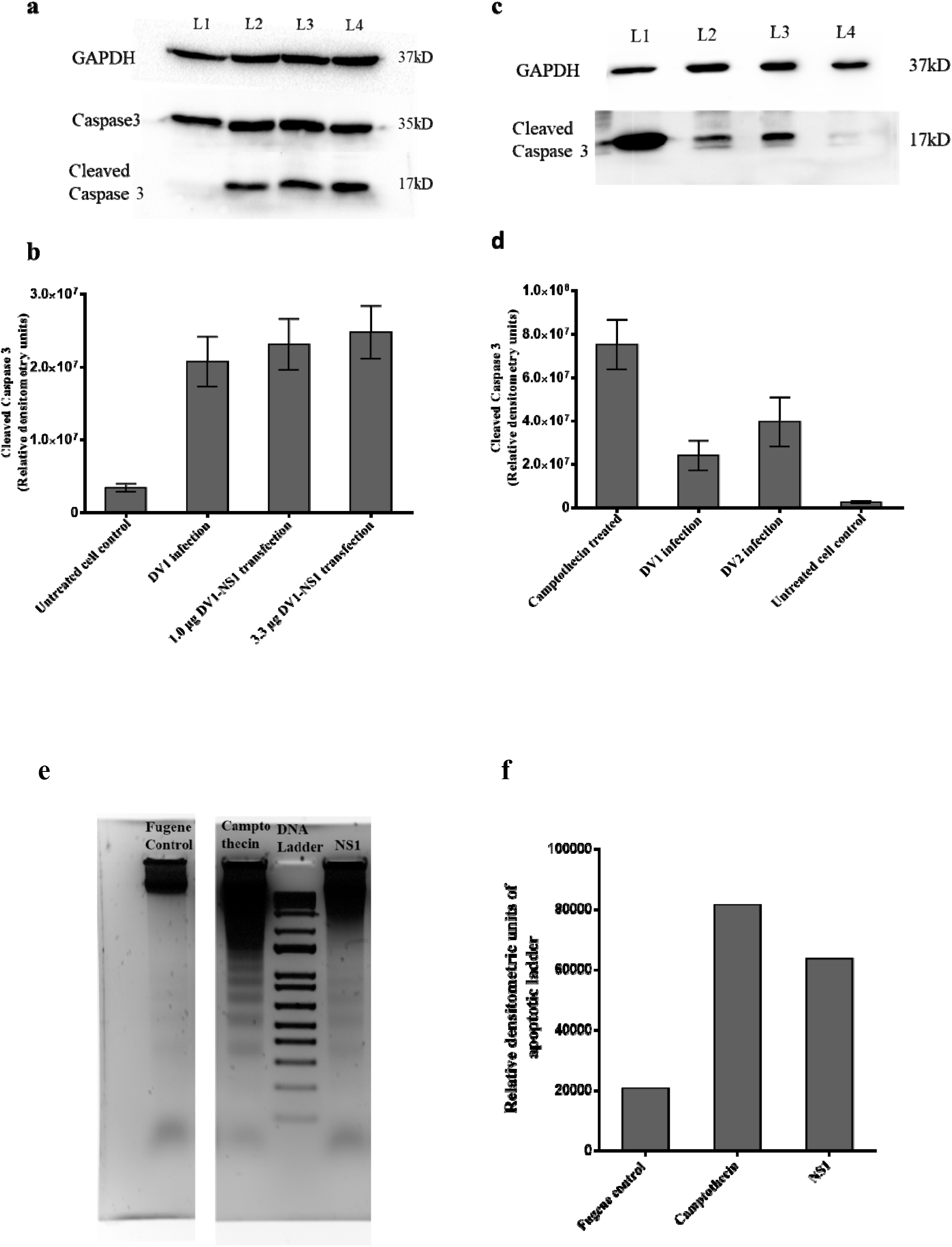
Dengue virus and NS1 only, both can induce Cleaved Caspase3. (**a**) Western blots data of Caspase 3 and Cleaved Caspase3 (CC3) expression in monolayer of Huh7 cells. Lane1: Untreated cell control; Lane2: DV1 infected cells (10 virus gE/cell); L3: 1.0 μg DV1-NS1-plasmid construct transfection; L4: 3.3 μg DV1-NS1-plasmid construct transfection. (**b**) Densitometry of bands from (a) for CC3. (**c**) Western blots data of CC3 expression in monolayer of Vero cells. Lane1: Cells treated with Camptothecin (4.0 μM) for 12h; Lane2: DV1 infected cells (10 virus gE/cell); Lane3: DV2 infected cells (10 virus gE/cell) R1; Lane4: Cell Control. (**d**) Densitometry of bands from (c) for CC3. GAPDH is as loading control. Both graphs show the average quantification from three independent experiments and error bars indicate SD. (e) Gel Electrophoresis image, from left, Lane 1 contains only Fugene treated Vero cells’ DNA; Lane 2 contains whole genomic DNA of Camptothecin (4.0 μM) treated Vero cells; Lane 3 contains DNA molecular weight marker; Lane 4 contains genomic DNA of DV1-NS1 (1.0 μg plasmid) transfected Vero cells. Equal amount of DNA was loaded in each well (f) Relative densitometric analysis of lanes of gel image as shown in (e).

DV1-HNSB-P4 infected Vero cells expressed less NS1 compared to Huh7, 0.4 (± 0.09) μg/ml at 62h post infection. Still such cells also showed higher CC3 levels compared to uninfected cells (Fig. 1c, d). So, NS1 alone can induce apoptosis as observed from increased expression of CC3 over cell control in transfection experiments. CC3 expression is considered a hallmark of apoptosis progression, so infected cells were supposed to undergo apoptotic DNA breaks. TUNEL and apoptotic DNA ladder experiments were performed to confirm this.

### Dengue virus shows control over apoptotic DNA breaks in DV infected cells

DV1-NS1 plasmid transfected Huh7 cells showed high percentage of TUNEL positivity, 29.17 (± 2.48) % for 1.0 μg plasmid and 36.67(±2.26) % for 3.3 μg plasmid. Increase in percentage of DNA nicks was positively linked to increasing plasmid transfection, confirming apoptosis induction by NS1. Surprisingly, DV1-HNSB-P4 infected cells, expressing similar concentration of sNS1 2.53 (± 0.79) μg/ml like 1.0 μg NS1-plasmid transfected cells (2.55 (± 0.59) μg/ml), showed insignificant levels of apoptotic DNA nicks 0.97 (± 0.32) %. In fact, the infected cells behaved very similar to untreated cell and mock transfection controls, 2.1 (± 0.40) and 1.5 ± (0.8) % respectively (Fig. 2a, b). So, DV1-HNSB-P4 infected liver cells expressed similar level of CC3 compared to NS1-transfected cells (Fig. 1a, b) but no subsequent DNA breakage was observed even up to 96h post-infection (Fig. 2a, b). This observation suggests that DV counteracts NS1 mediated-apoptosis of infected cells, as evident from the absence of DNA breaks.

**Fig 2.**
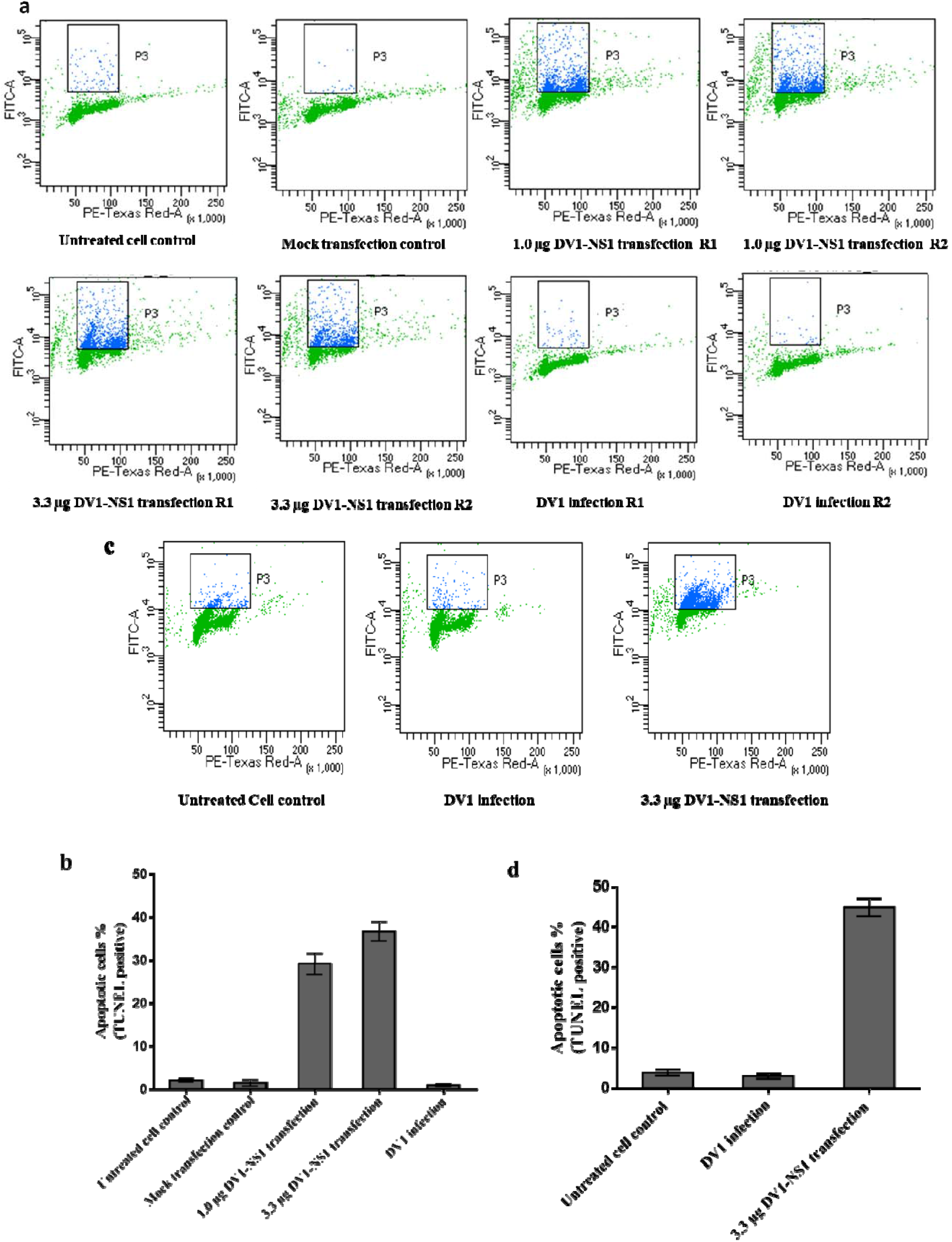
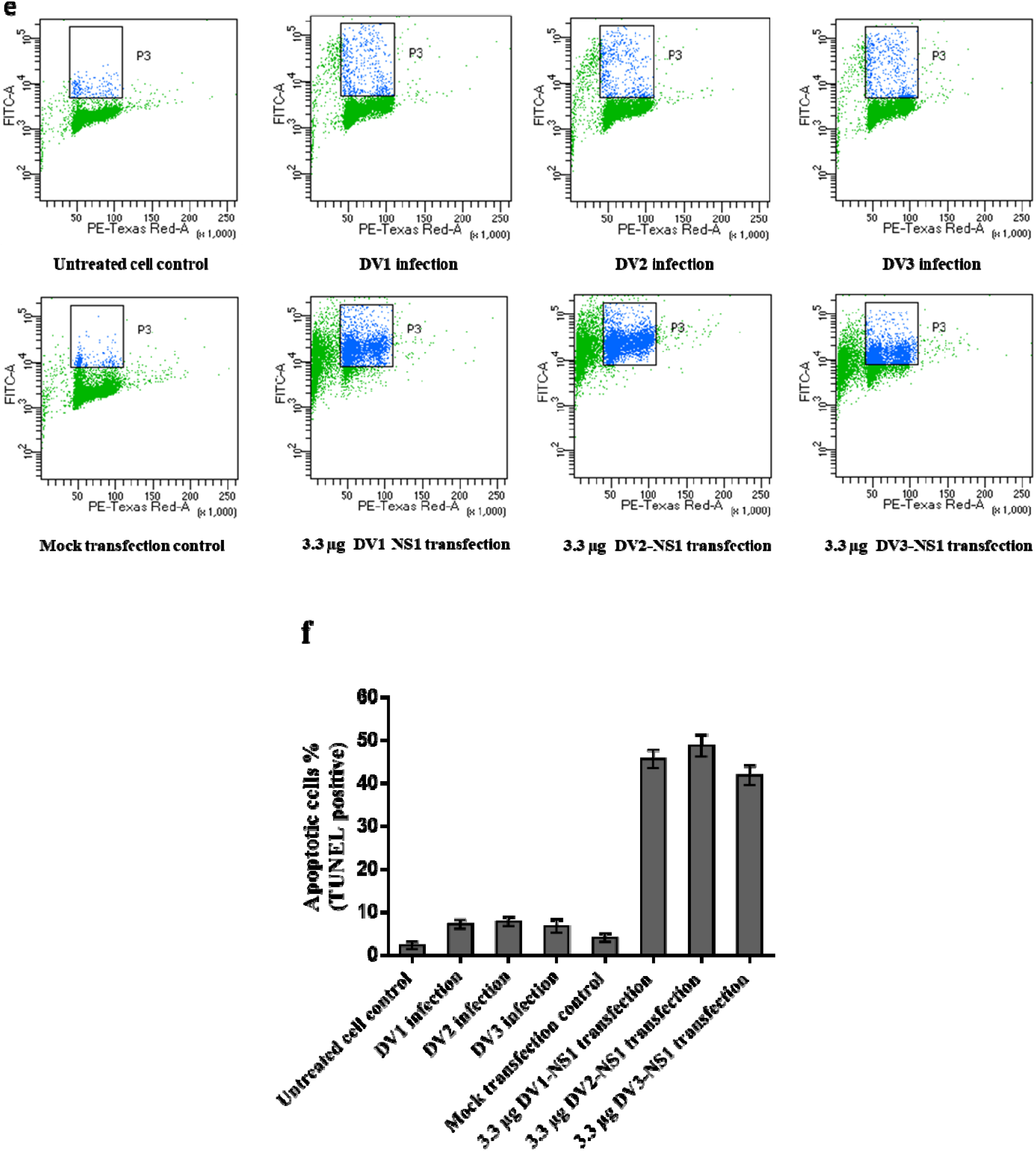
Dengue virus shows control/check over apoptotic DNA breaks in DV infected cells. **(a**) TUNEL assay representative data from monolayer of Huh7 cells, infected with DV1-HNSB-P4 (10 virus gE/cell) or transfected with 1.0 μg or 3.3 μg pcDNA3.1 (+)-DV1-NS1 construct/well of six well plate (as mentioned in figures). Cells were processed for TUNEL assay after 96h post infection or transfection. (**b**) Column graph of percentage of differentially treated cells that were found to be positive in TUNEL assay as in (**a**). (**c**) TUNEL assay data from monolayer of Vero cells with DV1 infection (10virus copies/cell) and transfection with 3.3 μg pcDNA3.1 (+)-DV1-NS1 construct (as mentioned in figures). (**d**) Column graph of percentage of differentially treated cells that were found to be positive in TUNEL assay as in (**c**). (**e**) Representative result of TUNEL assay from monolayer of Huh7 cells infected with 10 virus gE (DV1, DV2 or DV3) /cell, and cells transfected with DV1-HNSB-P4-NS1, DV2-HNSB-P4-NS1 or DV3-HNSB-P4-NS1 plasmid construct (3.3 μg) in each well of six well plate. (**f**) Column graph of TUNEL positive percentage of cells as in (**e**). Y axis of scatter plots (a, c, e) represents DNA breaks (BrdU labelling) and X axis represents DNA content (Propidium iodide staining). Column graphs show the average quantification from three replicates per condition and error bars indicate SD.

Similar study was also conducted multiple times using Vero cells. Again, there were more apoptotic DNA nicks in case of NS1 transfected Vero cells, confirming that only NS1 expression in cells was enough to induce apoptosis. DV1-HNSB-P4-infected Vero cells showed minimum TUNEL positivity (Fig. 2c, d) as observed in case of Huh7 cells.

The aforesaid phenomenon was found true for both DV2 and DV3. In case of DV2 clinical isolate, DV2-HNSB-NS1-transfected cells showed more apoptotic DNA breaks than only virus infected cells (10 virus gE/cell). 3.3μg DV2-NS1 plasmid transfected cells were more TUNEL positive (48.77 ± 2.55 %) than infected cells (7.83 ± 1.07 %) (Fig. 2e, f). Interestingly, this was observed with even higher levels of sNS1 production by DV2 infected (2.71 ± 0.08 μg/ml) than NS1-transfected cells (1.94 ± 0.11 μg/ml) (Supplementary Table1). So, despite higher level of NS1 expression in infected cells, apoptotic DNA breaks were lower compared to DV2-NS1 transfected cells. In case of DV3, the concentration of sNS1 in transfected cell supernatant was 1.56 (± 0.16) μg/ml in case of 3.3 μg DV3-HNSB-NS1 plasmid transfection. DV3-HNSB-P4 infected cells secreted 1.21 (± 0.15) μg/ml sNS1 (Supplementary Table1). Again, for DV3, NS1 expression in transfected cells resulted in 41.87 (±2.32) % apoptotic DNA breaks, whereas DV3 infected cells expressing equivalent concentration of sNS1 showed only 6.77 (± 1.55) % TUNEL positive (apoptotic) cells (Fig. 2e, f).

### Dengue virus exerts inhibitory effect over Camptothecin induced apoptotic DNA breaks in infected cells

Ladder assay was performed on repeated occasions with Camptothecin (4.0 μM) treated Vero cells as positive control. Camptothecin treated cells showed distinct ladder pattern resulting from DNA fragmentation which occurs at late apoptosis (Fig. 3a). NS1 transfected (1.0 μg) Vero cells’ DNA also showed apoptotic DNA ladder being consistent with CC3 expression (Fig 1e). In this experiment, DV1-HNSB-P4 infected cells were also treated with Camptothecin (4.0 μM) to promote apoptotic DNA breaks. Again, DV1-HNSB-P4 infected cells did not show apoptotic DNA ladder pattern even at 62h post-infection (point of cell harvesting), supporting the TUNEL assay results. Replication of virus was confirmed by 5-fold increase in sNS1 in supernatant over inoculum (Supplementary Table1).

**Fig 3.**
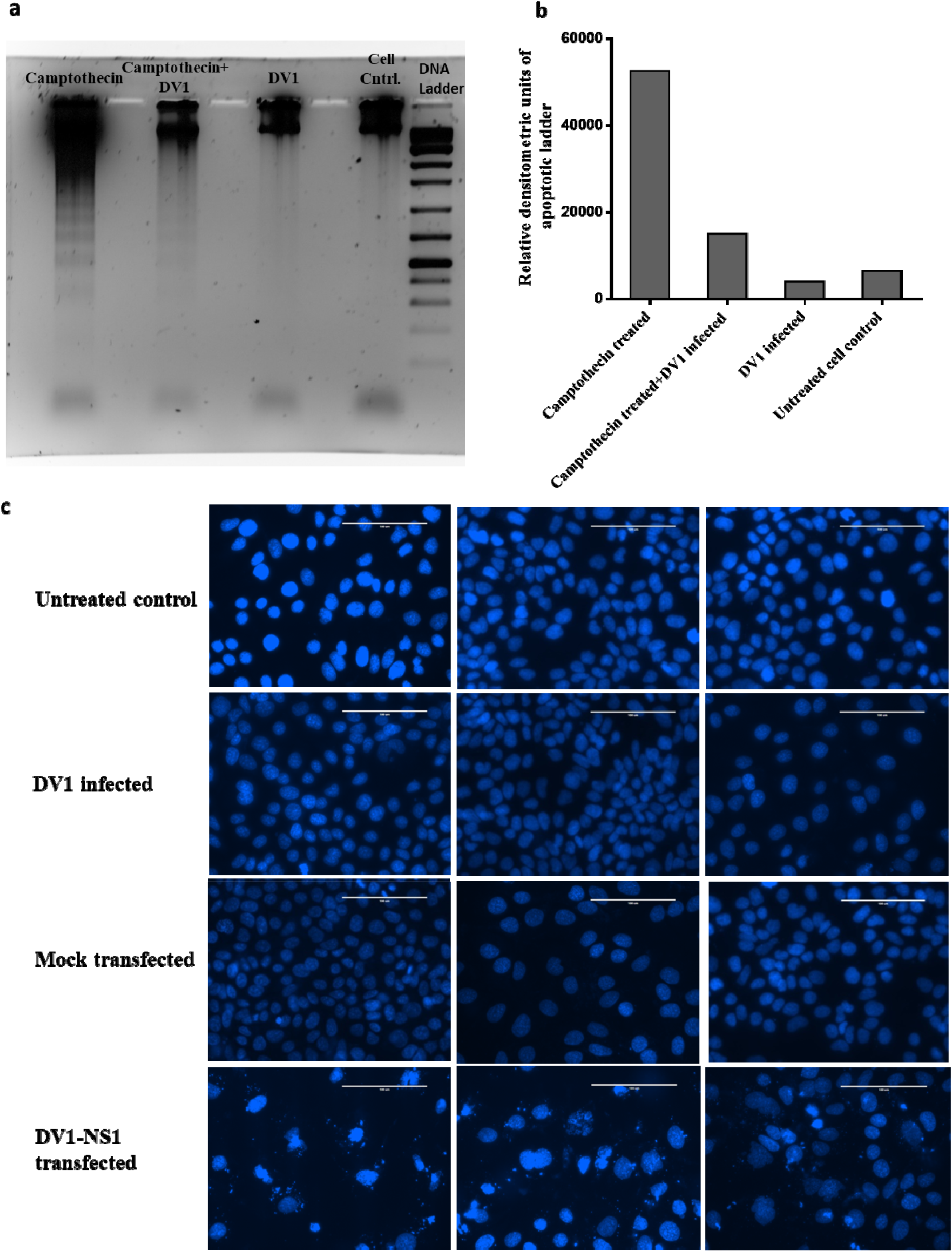
Dengue virus exerts inhibitory effect over induced apoptotic DNA breaks and counterbalances NS1 mediated nuclear damage. Monolayer of Vero cells were infected with DV1 at 10gE/cell. At 50h post-infection, one set of infected cells was treated with 4.0 μM Camptothecin for 12h, along with a control set of cells (no infection given) which was treated with only Camptothecin. At 62h post infection, all cells were harvested (**a**) Gel Electrophoresis image, from left, Lane1 contains whole genomic DNA of Camptothecin (4.0 μM) treated Vero cells; Lane2 contains DNA from Camptothecin (4.0 μM) treated DV1-HNSB-P4 infected (10 virus gE/cell) Vero cells; Lane3 contains DNA of only DV1-HNSB-P4 infected cells; Lane 4 contains DNA from untreated cells. Equal amount of DNA was loaded in each well. Lane5 contains DNA molecular weight marker. (**b**) Relative densitometric analysis of lanes of gel image (**a**). (**c**) Monolayer of Huh7 cells, grown on 22 mm coverslip were infected with DV1-HNSB-P4 (10 virus gE/cell). For transfection, 1.0 μg of plasmid containing the DV1-NS1 construct was used. 1.0 μg plasmid was chosen to keep the secreted sNS1 level similar to that in case of infected ones. In case of mock transfection, only Fugene was used without plasmid. Nuclei were stained with DAPI at 48h post infection or transfection. Images (40X) are representatives from multiple replicates from two different experiments. Scale bar, 100 μm. DV1: DV1-HNSB-P4, Cell Cntrl.-Untreated Cell Control.

DV1-HNSB-P4 infected cells together with Camptothecin (4.0 μM) treatment produced relatively reduced ladder pattern in comparison to only Camptothecin treated cells (Fig. 3 a, b). So, this competitive assay showed that Dengue virus is actually opposing apoptosis (by Camptothecin) and therefore, protecting the cellular DNA from fragmentation. This data conclusively proved that DV prevents cellular DNA breakage which is considered a salient feature of late apoptosis. Furthermore, it is observed that DV protects cellular DNA from damage even in presence of Camptothecin i.e. under high levels of CC3, as previously documented (Fig. 1c, d).

DAPI staining revealed that nuclear morphology of DV1 infected Huh7 cells was quite intact and similar to that of uninfected cells. In contrast, 1.0 μg DV1 NS1 plasmid transfected cells showed signs of nuclear damage in comparison to mock transfected (Fugene HD only) cells (Fig. 3c). We have already demonstrated that 1.0 μg DV1 NS1 plasmid transfected and DV1 infected (10 virus gE/cell) cells secrete comparable amounts of sNS1. It is therefore, expected that there should be similar levels of damage of nuclear morphology for both conditions. But nuclear morphology of DV1-infected cells was just like that of untreated cells, again providing direct visual evidence of Dengue virus’s control/check over apoptotic DNA damage and cytotoxic effects of NS1.

### Dengue virus keeps the infected host cell metabolically active reducing sNS1-mediated cytotoxicity

Metabolic activity of DV infected Huh7 cells (DV1, 2, 3) and their respective NS1 transfected cells were tested using MTT assay. Cells infected with DV at 10 virus gE/cell showed 89% viability in case of DV1; 86% with DV2 and 80% with the DV3 clinical isolate at 48h post-infection. Parallel sets of cells were transfected with respective NS1 constructs in amounts estimated to secrete equivalent amount of sNS1. It was observed that cell viability at 48h post-transfection was lower by 10-20% compared to DV infected cells, supporting the previously observed protective role of virus infection over NS1 cytotoxicity (Table 1).

**Table 1.** Percentage cell viability or metabolic activity as observed in MTT assay. Metabolic activities of infected cells were compared with uninfected controls. For NS1 transfected cells, metabolic activity was compared with mock Fugene HD transfected cells. For each type of treatment, there were four replicates, denoted as R1, R2, R3 and R4.

| **% of viable cells in infected cells**<br>**(10 virus gE/cell)** | **R1** | **R2** | **R3** | **R4** | **Mean (SD)** |
| --- | --- | --- | --- | --- | --- |
| DV1 | 73.57 | 110.08 | 80.79 | 91.34 | 88.95 (15.87) |
| DV2 | 90.94 | 90.55 | 81.77 | 82.35 | 86.40 (5.02) |
| DV3 | 89.97 | 67.71 | 87.23 | 73.37 | 79.57 (10.74) |
| **% of viable cells in NS1 plasmid**<br>**(0.11µg) transfected cells** | **R1** | **R2** | **R3** | **R4** | **Mean (SD)** |
| DV1-NS1 | 67.11 | 62.44 | 73.33 | 57.76 | 65.16 (6.65) |
| DV2-NS1 | 63.65 | 72.30 | 84.49 | 72.64 | 73.27 (8.56) |
| DV3-NS1 | 69.87 | 70.91 | 67.63 | 76.71 | 71.28 (3.87) |

### Release of mature virions as an evidence of successful virus replication in cell culture experiments

Secreted NS1 levels in experiments were measured using quantitative NS1 ELISA as described in methods. Individual ELISA, with standards, was performed for each experiment. In case of infection with DV1-HNSB-P4, DV2-HNSB-P4 and DV3-HNSB-P4, increase in sNS1 was expressed in microgram and as fold-increase over inoculum- sNS1 level (Supplementary Table 1). Increase in sNS1 level was one evidence of successful viral replication and NS1 production during every experiment. Interestingly, Camptothecin treatment did not have any significant effect on sNS1 production for virus-infected cells (Supplementary Table1).

Under same experimental conditions, Huh7 cells in six well plates were infected with the aforesaid three DV serotypes at 10 virus gE/cell. Inoculum was removed after virus adsorption and cell monolayer was washed with 1XPBS three times as done in case of all infections. Infected cells were harvested at 96h post infection and RNA was extracted. RNA from infected cell supernatant was also extracted to find the DV titer in it. Virus copy number was determined in cells and supernatant, using one step qRT-PCR(SYBR) and specific band size was confirmed in gel electrophoresis.

The means of intracellular virus copies were 4×10^7^ (±1.6×10^7^), 6×10^6^ (±7×10^4^) and 2×10^7^ (±10^6^) for DV1, DV2 and DV3 respectively. Similarly, supernatants of those infected cells also showed high virus titers, suggesting successful virus replication and release of mature virions. Mean total virus titers in DV1, DV2 and DV3 infected cell supernatants (3ml) were 4×10^7^ (±2×10^7^), 7×10^6^ (±5×10^6^) and 3.5×10^7^ (±10^7^) respectively. So, infected cells were secreting equal or more virus than intracellular titers (Table 2). The supernatant from the infected cells had been successfully used to infect fresh monolayer of cells, suggesting that the released virions in the supernatant were infectious, confirming their maturity and infectivity (data not shown).

**Table 2.** Virus copy number by qRT-PCR of RNA, extracted from infected Huh7 cells and supernatants.

| | Intracellular virus copies<br>Mean (±SD) | Virus copies in total<br>supernatant (3ml)<br>Mean (±SD) | Total yield of virus copies<br>Mean (±SD) |
| --- | --- | --- | --- |
| **DV1** | $4 \times 10^7 (\pm 1.6 \times 10^7)$ | $4 \times 10^7 (\pm 2 \times 10^7)$ | $8 \times 10^7 (\pm 4 \times 10^7)$ |
| **DV2** | $6 \times 10^6 (\pm 7 \times 10^4)$ | $7 \times 10^6 (\pm 5 \times 10^6)$ | $1.3 \times 10^7 (\pm 5 \times 10^6)$ |
| **DV3** | $2 \times 10^7 (\pm 10^6)$ | $3.5 \times 10^7 (\pm 10^7)$ | $5.5 \times 10^7 (\pm 10^7)$ |

**Table 3.**
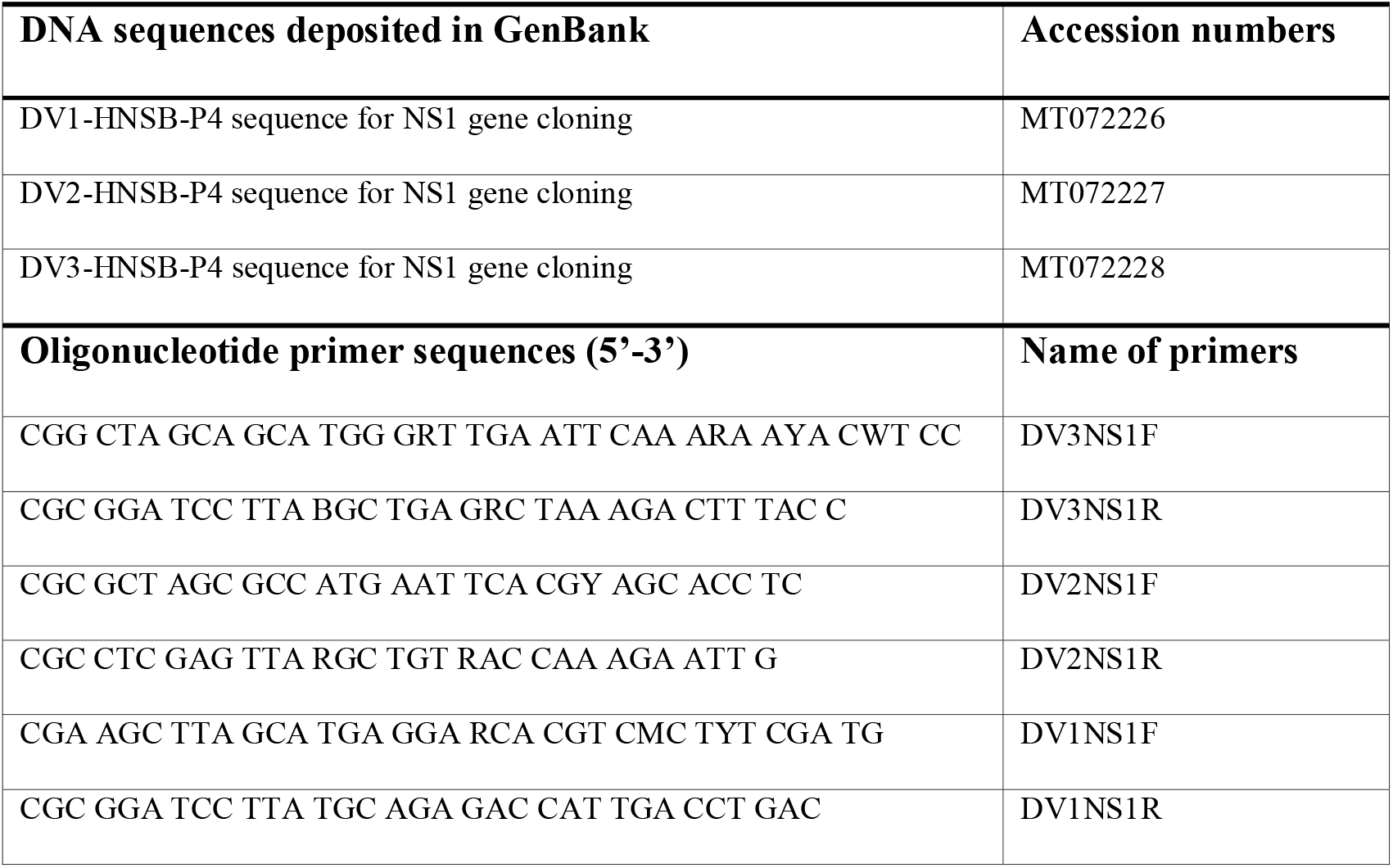
Dengue virus NS1 cloning primers and accession numbers of sequences.

To further confirm virus replication, immunofluorescence of DV infected Huh7 cells was done using anti-DV envelope primary Ab (DE1, Abcam) and goat anti-mouse IgG (Alexa fluor 488) secondary Ab. The presence of high virus titer was supported by viral envelope staining in more than 70% Huh7 cells. Nuclei of heavily infected cells were quite intact, again supporting DV-induced cell survival for increased virus replication (Fig. 4), opposing the deleterious effects of NS1 production and secretion. Primary Ab was developed based on envelope glycoprotein sequence of DV2 (Strain-16680) and worked best for DV2 staining, then DV1 but was found not effective for DV3 staining.

**Fig 4.**
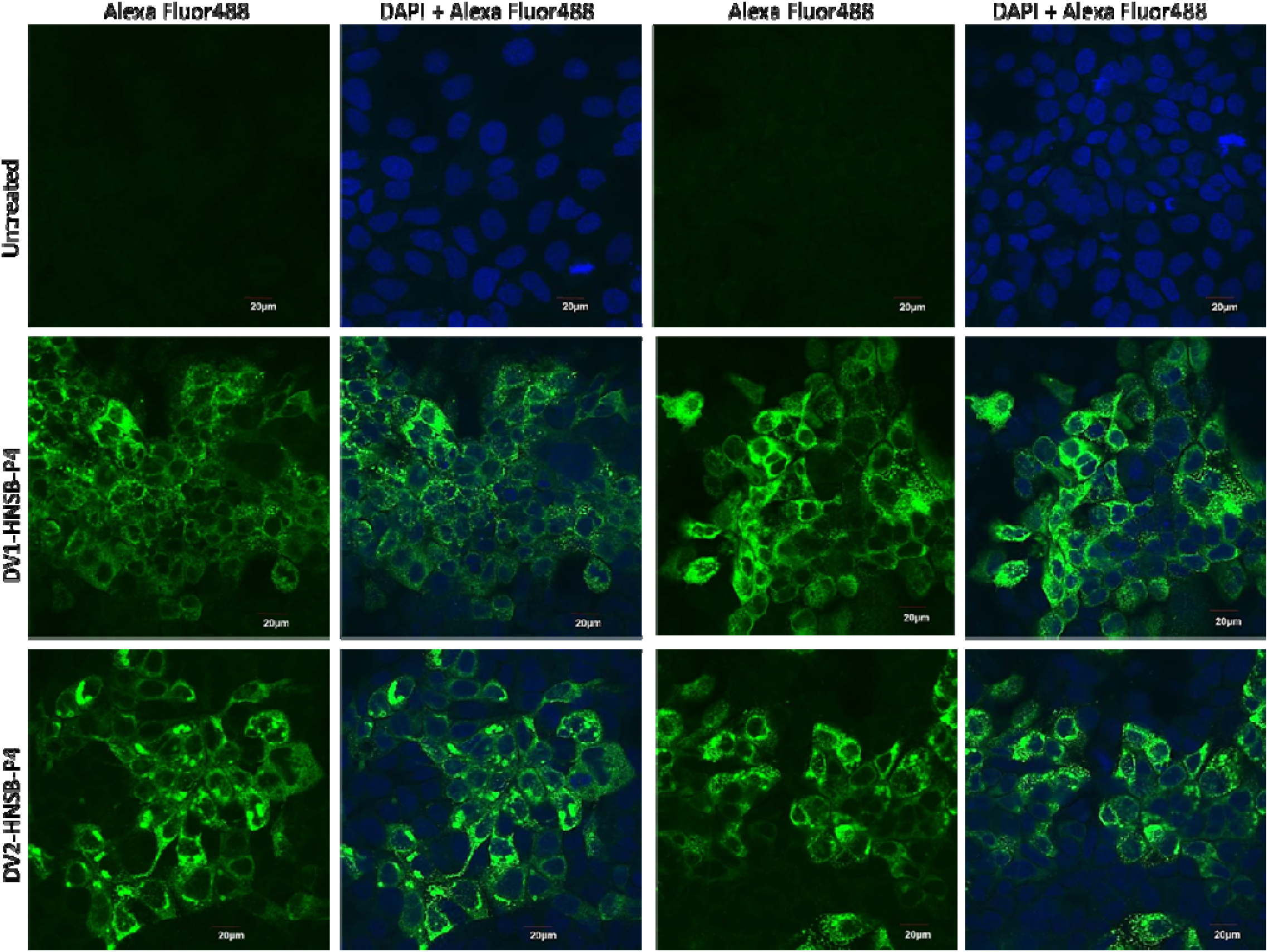
Immunofluorescence of viral envelope in DV1 or DV2 infected Huh7 cells. Monolayer of Huh7 cells was infected with DV1-HNSB-P4 or DV2-HNSB-P4 (10 virus gE/cell). At 96h post infection cells were fixed and treated with anti-DV envelope Ab and Alexa fluor 488 tagged secondary Ab. Nuclei of cells were stained with DAPI. Images (60X) are representative of multiple replicates from two different experiments. Scale bar, 20 μm. Two sets of images had been presented row-wise for each condition, each set comprising of again two images. For each set, one image shows only Alexa Fluor 488 staining (left one) and the other image of the same view, with DAPI and Alexa Fluor 488 (right one).

## Discussion

We have ascertained that Dengue virus-infected cells (Huh7 and Vero) and only NS1 gene-transfected cells, expressing equivalent amounts of sNS1 in the culture supernatant, both can induce cleaved caspase 3 (CC3). In case of transfection, sNS1 accumulation in the supernatant could not be due to a small fraction of cells as transfection with pcDNA3 EGFP (in similar amount to NS1-pcDNA3 plasmid) under identical conditions showed 60-70% cells to express GFP (Supplementary Fig. 1). This was also confirmed for NS1-pcDNA3 construct by FACS analysis (Fig. 2f). In case of transfection or virus infection (Fig. 4) (at 10 virus gE/cell), 60-70% cells were involved; both conditions produced around 2-3 μg/ml NS1 in the supernatant by 96 h post-infection which is comparable to physiological NS1 concentrations observed in clinical cases. In this study, we intended to examine NS1 at concentrations matching real case scenarios.

CC3 is a definitive marker of late apoptosis events that include subsequent fragmentation of cellular DNA. Our data suggest that virus-infected cells induced similar level of CC3 as observed for NS1-transfected cells (Fig. 1a, b). So apparently, NS1 contributes one major part in inducing apoptosis of DV-infected liver cells, the other part possibly comes from other pro-apoptotic factors like Capsid, Membrane, Envelope, NS2A and NS2B (18). The TUNEL assay results further confirmed that NS1 alone is capable of inducing apoptosis, resulting in DNA nicks, producing positive TUNEL signals (Fig. 2), apoptotic DNA ladder (Fig. 1e, f) and damage of nuclear morphology (Fig. 3c). In conclusion, our studies produced direct evidence that NS1 alone is capable of inducing apoptosis in the expressing liver cells at levels comparable to those found in real life DV clinical cases (Fig. 5) (14). We found no evidence that sNS1 (5.0 or 10.0 μg/ml) when transferred to fresh cell monolayers can induce apoptosis in such cells (tested at 96h post-transfer) by FACS analysis (Supplementary Fig. 2).

**Fig 5.**
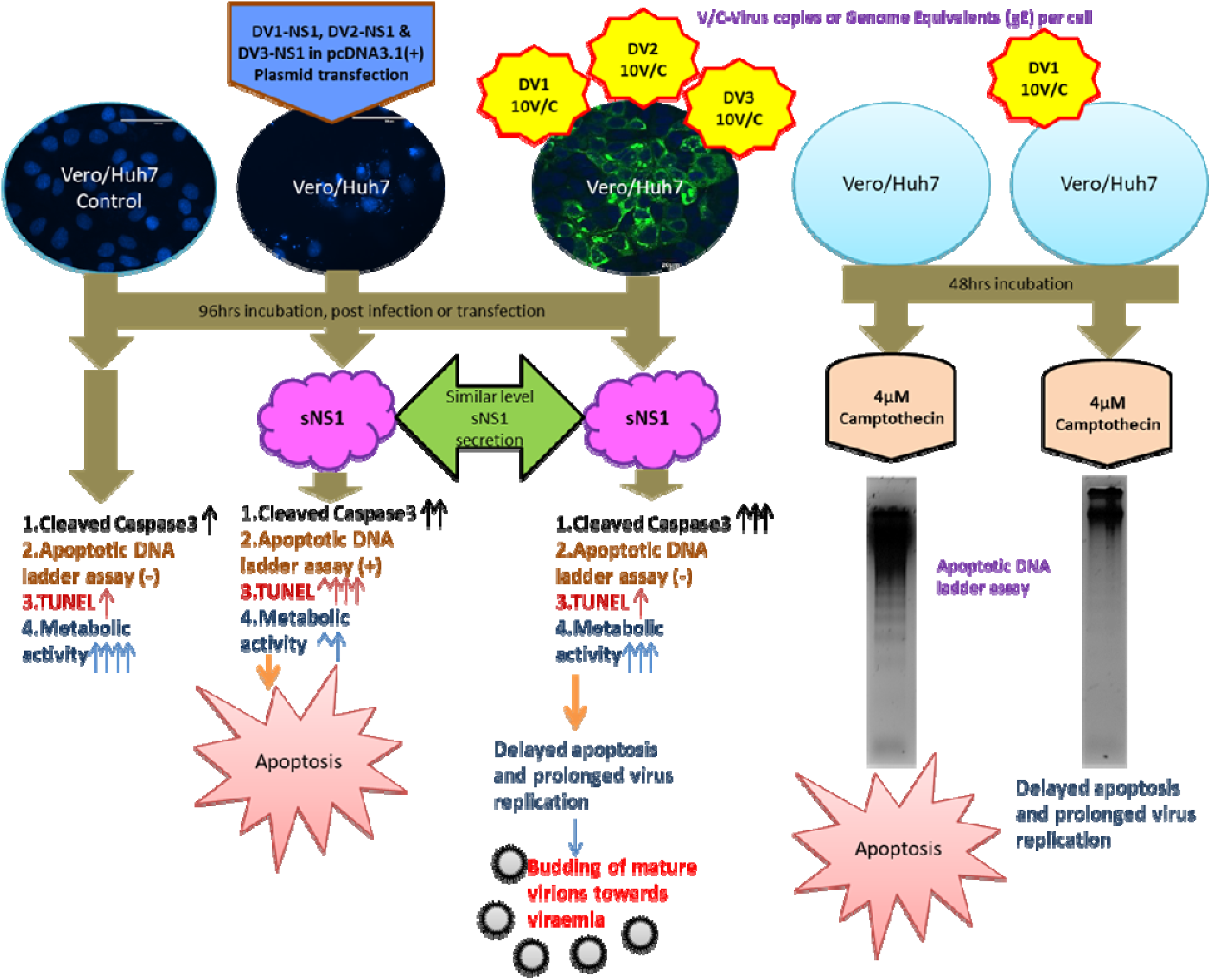
Dengue virus counteracts apoptosis in virus infected cells even in the presence of NS1 virotoxin that is alone capable of inducing apoptosis-schematic summary of the evidences.

It had been previously reported that DV infection in mouse neuroblastoma cell line, initiated survival signal via activation of PI3K and Akt phosphorylation. This survival signal was shown to prevent DV induced CC3 expression up to 24h (19). However, it is evident from our findings that DV infection actually results in CC3 expression, detectable even at 96h post infection. Possibly, PI3K-AKt activation-mediated survival signal appears to be transient lasting for only 24h post infection. It is apparent, that as DV infection progresses, CC3 is expressed to induce apoptosis plausibly to eliminate virus infected cells.

This obviously raised the question whether DV-infected cells really undergo usual apoptosis as part of host defence against viral infection? We found that although DV infection induces the apoptosis mechanism of host defence in infected cells, cellular DNA is protected from apoptotic damage and fragmentation (Fig. 2, 3). DV1 infected Huh7 cells expressed CC3 in similar levels as observed in case of 1.0 μg plasmid transfected cells. So, apoptotic DNA breaks as measured by TUNEL assay was supposed to be comparable in case of DV-infected cells and 1.0 μg plasmid-transfected cells. But it was found that virus-infected cells harboured much less apoptotic DNA breaks than transfected ones, in case of DV 1, 2 and 3 (Fig. 2a-f). This was further confirmed by the fact that DV infected cells, despite Camptothecin treatment, repeatedly showed reduced laddering pattern compared to only Camptothecin treated cells (Fig. 3a, b). Camptothecin is a Topoisomerase-I inhibitor and is known to induce apoptosis through DNA damage (20). It was therefore, apparent from our observations that DV infection counteracts apoptosis-mediated DNA breakage in Huh7 and Vero cells, even up to 96h post-infection. These point towards the ability of DV infection to delay the onset of apoptosis by preventing DNA breaks, besides inducing the early-stage host survival signal via PI3K-Akt pathway activation. The results of the MTT assays revealed that DV infection also keeps the host cell metabolically active counterbalancing host antiviral response i.e. apoptosis. On the other hand, NS1 expression and secretion from cells in absence of whole virus led to substantial metabolic damage or mortality of cells (Table 1).

There are several reports presenting evidences of DV-mediated apoptosis of liver cells ^(21–23)^. In these studies, experiments were carried out with laboratory strains of Dengue virus. Our studies involved three clinical isolates of Dengue virus, representing serotypes 1, 2 and 3, circulating in Kolkata during 2017. DV isolates with low passage number (passage number 4 for all three isolates) were used in all assays in order to maintain them closer to the wild-type circulating isolates and to mimic the clinical scenario as much as possible (24).

Our observations perhaps explain why majority (≥80%) of DV infections in humans are asymptomatic/self-limiting (25) despite the fact that sNS1 on its own is a virotoxin capable of causing cytotoxicity/apoptosis of liver cells. The results also prove that sNS1-mediated cytopathic effects are suppressed/ameliorated in case of virus-infected cells. These observations encouraged us to propose that DV disease outcome/severity may depend on the “tilt” of the balance between DV infection-mediated prolongation of cell viability (by check of cellular apoptosis and resulting DNA damage) and induction of apoptosis by proapoptotic virus proteins like NS1. In this context, it is interesting to note that symptomatic to severe DV cases have been implicated in case of the following conditions/pre-disposing factors, namely (I) Antibody dependent enhancement (ADE) in case of secondary infection (26); (II) Level of sNS1 and viraemia (14,27); case fatality was high when NS1 antigenaemia was high as observed in case of DV2 infections. In contrast, higher case fatality was linked to higher viraemia in case of DV3 infections (14,27); (III) Co-infection with different serotypes (28); (IV) Immune status or comorbidities of the host. It appears that the above factors decide the fate of DV infection affecting the aforesaid “tilt” of the balance between asymptomatic condition (self-limited recovery) and disease progression.

Virus replication in Huh7 and Vero cells was confirmed in infected cells. Alongside high intracellular virus titer, infected liver cells also secreted equivalent or more mature virions, suggestive of successful virus infection of cells resulting in high viraemia. Mature virions were released keeping the cells intact as evident from high virus copy numbers in infected cell supernatants (Table 2). Additionally, DV replication in individual experiments was confirmed by fold increase in sNS1 level in supernatants (Supplementary Table1).

It has been observed that clinical isolates of DV are not plaque-forming at early passages in cell (29). The same appears to hold true for our clinical isolates. Enveloped DV in infected cells was also visualized by immunofluorescence using Ab against DV envelope glycoprotein (Fig. 4). The nuclei of DV-infected Huh7 cells were similar to those of the uninfected cells and were in sharp contrast to the damaged nuclei in case of NS1-transfected cells (Fig.3c). This observation again emphasized DV mediated survival of infected host cells in the presence of NS1 virotoxin in the background.

Apoptotic DNA fragmentation is triggered by the nuclease DNA fragmentation factor 40 (DFF40). DFF40 expression and folding are regulated by DFF45. The latter acts as a nuclease inhibitor prior to DFF40 activation mediated by execution caspases like Caspase3. Activated execution caspases cleave DFF45 which then dissociate from DFF40, leaving DFF40 as the functional nuclease (30). The DFF40 and DFF45 expression levels had been checked by means of Western blot and no significant difference was observed between the infected and the transfected cells (data not shown). It is still not clear how DV exactly protects infected cells’ DNA. The explanation appears to be not straightforward and warrants further in-depth investigation, it may be due one or more viral factors or modulated host factors or a combination of both.

Our finding also corroborates well with the fact that Dengue virus infection is associated with increased cell free DNA (cfDNA) in plasma. There are several reports that cfDNA increases in DV infected patients and is associated with disease severity (17). In this article the authors have said cellular apoptosis to be the source of this high cfDNA, but the mechanism was not clear; as apoptosis was supposed to produce fragmented DNA, to be removed soon via salvage pathways and excretion. But if DV infection protects cellular DNA from apoptotic breaks, that will result in larger DNA fragments which may stay in plasma for longer period. Our results possibly explain this cfDNA in Dengue infections more rationally. Disease severity is positively linked to DV titres; more virus replication means infection of more cells of the body; more cells rupture to release progeny virions and “protected” cfDNA-this perhaps explains why more cfDNA in plasma has been recorded as a biomarker of DV disease severity (31).

DV infection of cells definitely induces apoptosis as a response of host defence (19). Adding to this, we have shown NS1 alone is capable of inducing apoptosis, besides other well-known pro-apoptotic factors like virus capsid protein. So, it is a viral strategy against host defence that DV infection “protects” cellular DNA from apoptosis-induced fragmentation, to keep the host cell metabolically active even with toxic NS1 expression as observed. This would allow the virus a window to replicate for a relatively longer period using the host cellular machinery (we recorded up to 96h post-infection) followed by budding of mature virions (Table 2). This strategy is advantageous from the point of DV infection biology.

In a nutshell, virus-mediated “protection” of cellular DNA allows more time and space for virus replication within infected cells, in the face of ensuing apoptosis, thereby allowing the virus to reach higher titres to establish successful viraemia in the host. Successful virus transmission and propagation in an endemic region occurs when the virus is present in high titres in circulation so that the mosquito vectors during blood meal could take in adequate number of infectious virus particles, thereby ensuring effective virus transmission and persistence in the host and vector populations.

## Materials & methods

### Cell Lines

Vero and Huh7 cells were obtained from NCCS, India. Cells were cultured in DMEM (D5796, Sigma) supplemented with 10% FBS (Gibco) and Pen-Strep and L-Glutamine mix (Sigma). Monolayers of cells during experiments were maintained using DMEM supplemented with 1% FBS. Cells were grown at 37°C with 5%CO_2_. During passage, cells were washed with PBS (1X) and detached with Trypsin-EDTA (1X) (Gibco). For culture, Thermo-Nunc flasks and culture plates were used. C6/36 cells were cultured in MEM (Sigma), supplemented with 10% FBS (Gibco), Pen-Strep and L-Glutamine (Sigma) and Fungizone (Gibco). C6/36 cells were grown at 28°C with 5%CO_2_.

### Viruses

Serum samples from Dengue fever patients were collected and supplied by Dr. Keya Basu and Dr. Abhishek De with proper information and prior written patient consent. This study was approved by the Ethics Committee on Human Research of CSIR-Indian Institute of Chemical Biology, Kolkata. All serum samples were confirmed as Dengue virus-positive by means of NS1 diagnostic ELISA test (Platelia, Biorad). Serum samples were filtered using Millipore 0.22 μm PES syringe filters. 70% confluent monolayer C6/36 cells were infected with filtered serum (inoculum volume made up to 800μl in MEM for T-25 flask of C6/36 cells); adsorption was done for 2h under normal cell culture conditions with intermittent shaking at every 15minutes. Cells with virus were incubated for 120h. Three such passages were given in C6/36. During harvesting T-25 containing cells were scraped in 1ml supernatant. These cells in supernatant were freeze-thawed and centrifuged at 13,000 rpm for 15min at 4°C to pellet the cellular debris. The resultant clear supernatant was aliquoted and stored at −80°C as virus stocks. Virus serotyping was done as described by Lanciotti et al (32). Virus titer was determined using SYBR-based one step qRT-PCR with Luna Universal One Step qRT-PCR reagent (NEB). QuantStudio 5 (Applied Biosystems) was used to run the qPCRs. Primers as described by Lanciotti were used in qRT-PCRs.

### TUNEL assay

TUNEL assay was done as per protocol of APO-BrdU™ TUNEL Assay Kit with Alexa Fluor™ 488 Anti-BrdU (Thermo, Life Tech). Adherent cells were trypsinized and pelleted along with floating cells. The pellet was resuspended in 600μl 1X PBS. Then 4.4 ml of 70% chilled ethanol was added, keeping the cells suspended by mild vortexing. Cells in 70% ethanol were stored at −20°C overnight. Next day, cells were centrifuged at 300g at 4°C for 5mins and the supernatant was decanted. Cells were then washed twice with 1ml wash buffer. Thereafter, the pelleted cells were suspended in 50 μl reaction mix and incubated at 37°C for 70mins. During incubation, tubes were tapped at every 15mins interval to keep the cells in suspension. After incubation cells were rinsed twice with 1ml of rinse buffer. Then 100 μl of antibody mix (5 μl in 145 μl rinse buffer) was added to each tube, keeping the cells suspended by tapping. Cells with antibody were incubated at room temperature (~25°C) for 30mins. The cells were then washed once with 700 μl rinse buffer. After that, cells were suspended in 500 μl Propidium iodide (PI)-RNaseA solution and transferred to FACS tube. Cells were allowed to incubate with PI for 30mins, followed by FACS analysis in BD LSRFortessa. Analysis of data was done using the BD FACSDiva 8.0.2 software. In TUNEL data analysis, gating for virus infected cells was based on uninfected cell control. In case of transfection experiments, gating was based on mock transfection control.

### Apoptotic DNA Ladder assay

Adherent cells in T-25 flasks were detached using trypsin-EDTA and pelleted by centrifugation. Pelleted cells were lysed using cell lysis buffer (1X PBS, 0.2% TritonX100). RNaseA (Invitrogen) was added (2 μl) to the cell lysate (about 200-300 μl) and the resultant mix was incubated at RT for 5mins. This was followed by addition of 20 μl Proteinase K and incubation at 56°C in heat block for 2h. Then equal volume of Phenol-Chloroform-Isoamyl alcohol (25:24:1) (HiMedia) was added and mixed by inverting. This was followed by centrifugation at 13,000rpm for 15mins. Of the three layers visible, upper most transparent layer was aspirated carefully to a new 1.5 ml Eppendorf tube. Isopropanol (1.2 times the volume of aspirated supernatant) was added to the supernatant and mixed well by inverting. 1 μl of Glycogen (Invitrogen) was added to the tube before storing it at −20°C for 2hr. The tube was then centrifuged at 13,000rpm for 15mins at 4°C to obtain the DNA pellet. Supernatant was discarded and pellet was washed using 70% ethanol twice. Pellet was air dried and suspended in 40μl nuclease-free (NF) water (Ambion). DNA quantity and quality were ascertained using Nanodrop One (Thermo). Equal quantities of cellular DNA from different conditions of the experiments were subjected to gel electrophoresis in 1.4% agarose gel with SYBR safe dye (Invitrogen). Agarose gel electrophoresis was done at 50V (5V/cm) for 3hr. Gel was observed in Gel Logic (Carestream) under UV transillumination.

### NS1 ELISA

ELISA was done as per protocol of Bio-Rad Platelia Dengue NS1 ELISA. In case of quantitative ELISA, serial dilutions of recombinant NS1 Antigen (Ag) (Bio-Rad) was used. Dilutions of Ag were made in PBS (1X). ELISA reading was taken in iMark plate reader (BioRad). Standard curve was generated from ODs of known dilutions of NS1-Ag. From the equation of that curve, soluble NS1 quantity of unknown samples were determined. Separate ELISAs were performed for each experiment.

### Western Blot experiments to detect Cleaved Caspase3

Monolayer of Vero or Huh7 cells was subjected to Camptothecin (4μM) treatment for 12h in case of NS1 transfection or Dengue virus infection. Lysis buffer (1% TritonX100 in PBS, 10U/ml DNase1-Cat. No. D2821, Sigma with Proteinase inhibitor (Pierce, Thermo)) was directly applied on adherent cells and kept on ice for 10mins. Lysate was aspirated into 1.5ml tube and centrifuged at 13,000rpm for 15mins at 4°C. Clear supernatant was aspirated and used for Western blotting. Protein quantification was done using BCA assay kit (Pierce). Cell Lysate was separated by means of 5% (Stacking) and 15% (Resolving) SDS PAGE gel electrophoresis, using running buffer (25mM Tris, 192mM Glycine, 0.1% SDS: pH 8.3) at constant voltage of 90V for 2.5h. To observe the relative position of protein in gel, two protein ladders, namely PageRuler (Thermo) and Precision Dual Colour ladder (Bio-Rad) were used. For subsequent analysis proteins were transferred onto Nitrocellulose membrane using Trans-Blot Turbo Transfer System (Bio-Rad). Membrane was blocked using TBST with 5% skimmed milk for two hours at room temperature. Anti-Caspase 3 Ab (CST-9662) (1:1000) and anti-cleaved Caspase 3 Ab (CST-9661) (1:1000) were used to detect the target proteins of interest. Primary anti GAPDPH Ab (CST-8884S), HRP conjugate (1:2000) was used as housekeeping protein control. Strips of membranes corresponding to target proteins (Caspase3, CC3 or GAPDPH) were incubated with Primary antibodies, overnight at 4°C with fresh blocking buffers. Secondary anti-rabbit IgG-HRP (Abcam-97051) (1:2000) was incubated for 1.5h with strips. The membranes were treated with ECL substrate (Bio-Rad) prior to visualization of the stained protein bands in Azure Biosystem Chemi Doc system or ChemiDoc (BioRad). Densitometric analysis was done using ImageJ and Image Lab software (Bio-rad).

### Cloning and expression of NS1 genes of Dengue virus types 1, 2, 3

RNA from virus was extracted using High Pure Viral Nucleic acid extraction kit (Roche). Using gene specific reverse primers of target genes, cDNAs were synthesized using Superscript-III (Invitrogen). Respective cDNA was used as template to PCR-amplify the desired target. PCR products were confirmed by agarose gel electrophoresis using 0.8-1% agarose and purified using PCR purification kit (Qiagen). In case of NS1 gene, for all three serotypes, the gene corresponding to signal sequences of 23-29 amino acids upstream of NS1 protein was included for NS1 gene amplification for cloning purposes. Primers for cloning contained appropriate restriction enzyme cutting sites, for instance, HindIII and BamHI restriction sites in forward and reverse primers respectively in case of DV1 NS1 gene (plus signal sequence). In case of DV2 NS1, NheI and XhoI restriction sites were incorporated in forward and reverse primers respectively. For, DV3 NS1, forward primer contained NheI and reverse primer contained BamHI restriction sites. Restriction enzymes (NEB) digestion was carried out for respective PCR products and the expression vector pcDNA3.1 (+). Ligation was done using T4 DNA ligase (Promega), followed by transformation in XL1-Blue cells. Plasmid was purified using Purelink-Midi plasmid purification kit (Invitrogen). Plasmid transfection was done using Fugene HD (Promega), following manufacturer’s instructions.

### MTT assay

Huh7 cells were seeded as monolayer in 96 wells plate. Cells were infected with DV or transfected with NS1 gene construct. One column of cells was left uninfected/un-transfected, as positive control while another column was not seeded with cells but filled with media only, as negative control. Media was removed 48h post infection/transfection, from each well and the cells were washed using 1X PBS. Then 90 μl DMEM (Phenol red-free) and 10 μl MTT solution were added to each well. The plate was incubated at 37°C and 5% CO_2_ for four hours in a humidified CO_2_ incubator for the formation of Formazan crystals in the living cells. Media was discarded from all wells; 100 μl DMSO was added to each well and the contents of the wells were mixed properly to dissolve the crystals. The plate was incubated for 30mins and absorbance was measured at 590nm wavelength.

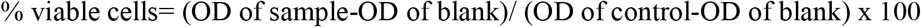

### Immunofluorescence experiments

For DAPI-staining, monolayer of Huh7 cells was cultured on 22mm cover slips and treated (DV infection or NS1 gene transfection) or left untreated as needed. Media was removed at 48h post-treatment and cells were fixed with ice-cold 70% ethanol. This was followed by one wash with nuclease free water (Ambion). Cover slips were then soaked using lint free paper and mounted with Prolong Diamond antifade mountant with DAPI (Invitrogen). Images were taken using EVOS FL cell imaging system.

In case of DV envelope immunofluorescence, monolayer of Huh7 cells was grown on six chambered slide (Genetix). DV1, DV2 or DV3 were inoculated at 10 virus gE/cell. Virus adsorption was done for 2h. At 96h post infection, media was removed and cells were washed twice with PBS. Cells were subsequently fixed using 4% PFA in PBS for 10min at RT. Cells were again washed twice with 1XPBS. Cells were permeabilized with 0.1% Triton X100 in 1XPBST for 15mins at RT. Then blocking was done with 1%BSA and 22.52mg/ml Glycine in PBST for 45mins at RT on rocker. Cells were incubated with diluted (1:40) primary Ab (Ab41349) in 1% BSA in PBST for one hour at RT on a rocking platform. The antibody solution was then removed by decanting and the cells were washed three times with PBST, each wash for 5mins. The cells were then incubated with diluted (1:500) secondary Ab (Ab150113) in 1% BSA in PBST for 30mins at RT. Secondary Ab was removed and cells were washed four times with PBST, each of five min duration, on rocking platform. The slides were allowed to get dry and mounted with Prolong Diamond antifade mountant with DAPI (Invitrogen). Images were taken using Fluoview 10i, Confocal microscope (Olympus).

## Acknowledgements

We thank Mr. Tanmoy Dolui for his advice during TUNEL assay. Thanks to Mr. Subrata Roy for his suggestions in wet lab work. H.N. thanks Mr. Tathagata Kayal for his help during cell culture. H.N. and A.G. acknowledge Council of Scientific and Industrial Research (CSIR) for Senior Research Fellowship (NET-SRF) for pursuing this work. This work was supported by CSIR-IICB institutional funding (MLP-118). The authors acknowledge CSIR-IICB for providing laboratory facilities for conducting the current work.

## Author Contributions

S.B. and H.N. conceived and designed the experiments. K.B. and A.D. collected and supplied the clinical samples. H.N. performed majority of the experiments. A.G. also performed many of the experiments. S.B., H.N. and A.G. performed data analysis and S.B. performed critical analysis of the data. S.B. provided funding, reagents, materials and analysis tools. S.B. and H.N. jointly wrote the initial drafts. All authors critically reviewed and modified and agreed on the current version of the manuscript.

## Ethical approval

This study was performed in accordance with the ethical standards (at par with the 1964 Helsinki declaration and its later amendments) of the review boards of all relevant institutions. Ethical approval for the research was granted by the respective Institutional Ethical Committee of CSIR-IICB and Calcutta National Medical College, Kolkata. All experiments were carried out in accordance with the relevant guidelines and regulations.

## Declaration of Interests

The authors declare no competing interests.

## Notes

### Competing Interest Statement

The authors have declared no competing interest.

### Summary of Updates

Several sections including the Supplemental files have been updated.

