## Supplementary Table1 for "Dengue virus clinical isolates sustain viability of infected hepatic cells by counteracting apoptosis-mediated DNA breakage"

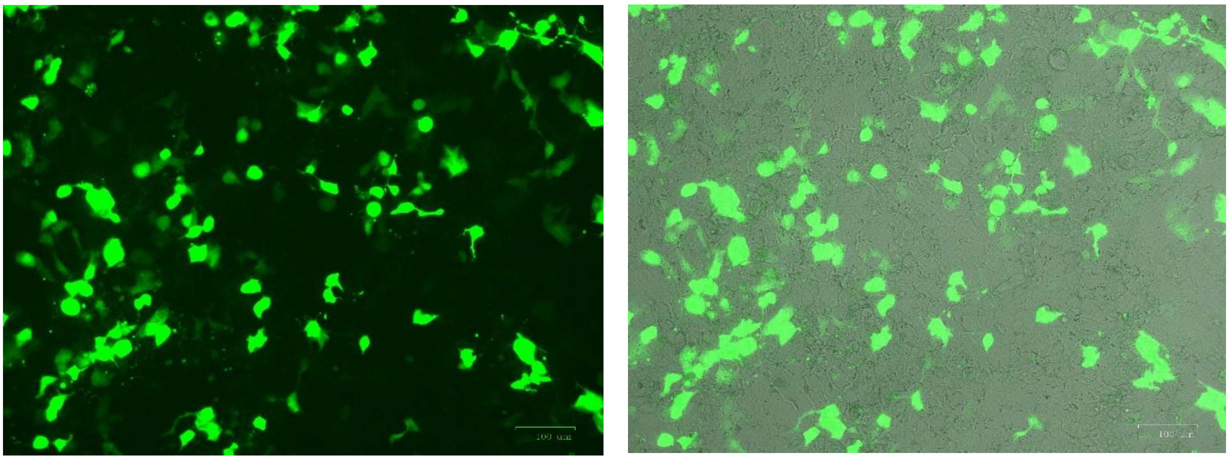

**Fig. S1. Monolayer of Huh7 cells transfected with pcDNA3-EGFP plasmid**

Huh7 cells were transfected with 1.0 µg pcDNA3-EGFP plasmid using Fugene HD. Images were taken at 48h post-transfection. Left panel shows only GFP expressing cells. The right panel shows merged image of phase contrast view of cells including the GFP expressing cells. Scale bar 100 µm.

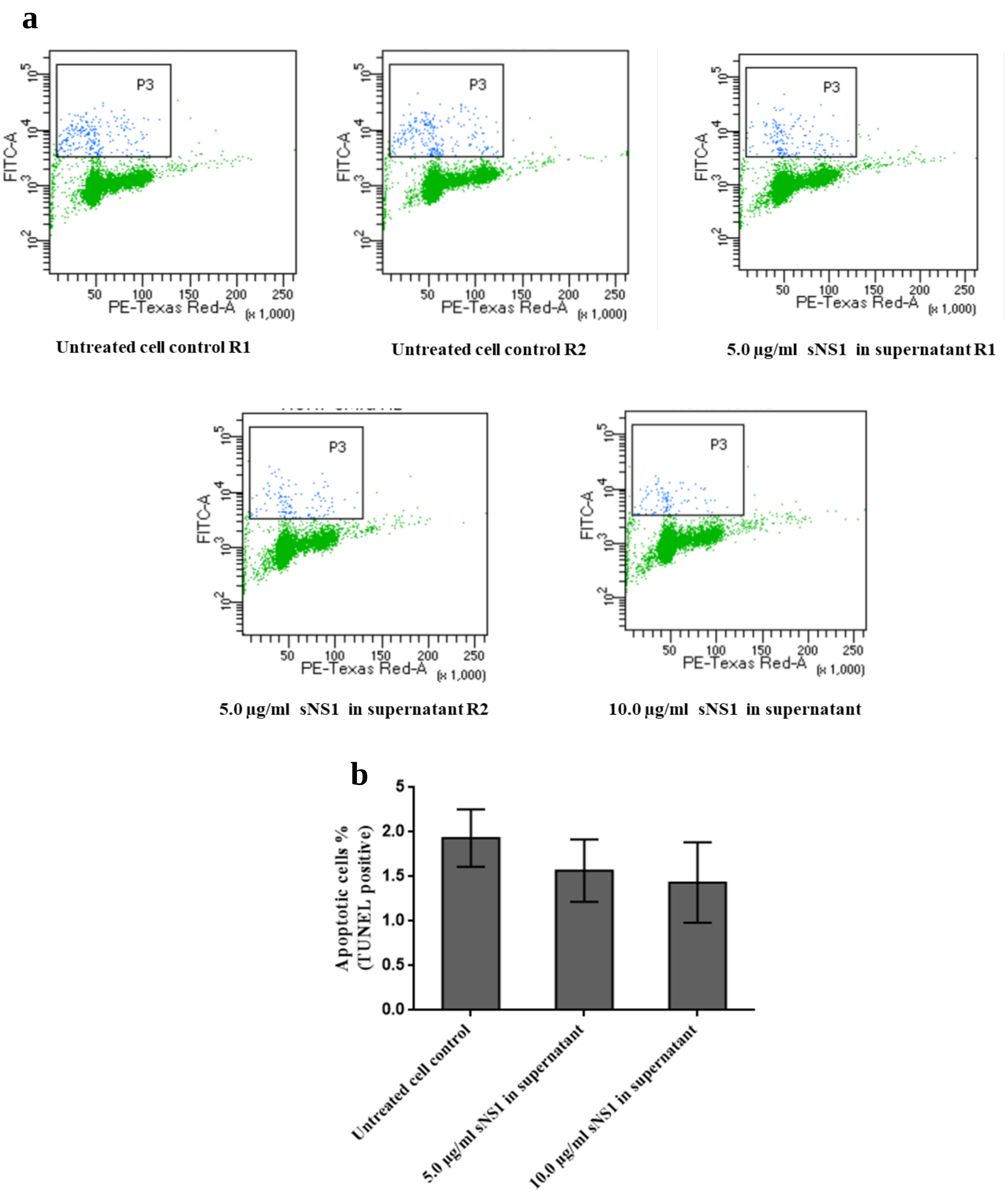


**Fig. S2. Soluble NS1 (sNS1) in supernatant of fresh Huh7 cells, could not induce apoptosis**

(**a**) Representative TUNEL assay data from experiments where monolayer of Huh7 cells in six well plates were treated with 5.0 µg/ml or 10 µg/ml sNS1 in supernatant for 96h. (**b**) Column graph of percentage of sNS1 treated cells that were found to be positive in TUNEL assay as in (**a**). Y axis of scatter plots (**a**) represents DNA breaks (BrdU labelling) and X axis represents DNA content (Propidium iodide staining). Column graphs show the average quantification from three replicates per condition and error bars indicate SD.

| Experiment with relevant Figure number | Type of cells & type of culture plate | Sample identification | Infection of DENV/ Transfection of NS1 | NS1 in Inocu-lum (µg) | Yield of NS1 (µg) excluding NS1 in inoculum | Fold change of NS1 (µg) over inoculum | Yield NS1 conc. (µg/ml) | Time of harvesting after infection or Transfection | Virus gE/cell | Transfected plasmid (µg) | Mean yield (µg/ml) | SD |
|---|---|---|---|---|---|---|---|---|---|---|---|---|
| Fig:1(a) WB | Huh7, 6well | DV1-HNSB-P4(R1) | Infection of DV1-HNSB-P4 | 0.48 | 10.23 | 21.26 | 3.41 | 96hrs post infection | 10 | NA | **2.53** | **0.79** |
| | Huh7, 6well | DV1-HNSB-P4(R2) | Infection of DV1-HNSB-P4 | 0.48 | 8.71 | 18.10 | 2.90 | 96hrs post infection | 10 | NA | | |
| | Huh7, 6well | DV1-HNSB-P4-NS1(R1) | Transfection of DV1-HNSB-P4-NS1 | NA | 8.9 | NA | 2.97 | 96hrs post transfection | NA | 1 | **2.55** | **0.59** |
| | Huh7, 6well | DV1-HNSB-P4-NS1(R2) | Transfection of DV1-HNSB-P4-NS1 | NA | 9.35 | NA | 3.12 | 96hrs post transfection | NA | 1 | | |
| | Huh7, 6well | DV1-HNSB-P4-NS1(3.3) | Transfection of DV1-HNSB-P4-NS1 | NA | 14.22 | NA | 4.74 | 96hrs post transfection | NA | 3.3 | **4.36** | **0.33** |
| Fig2:(a) TUNEL | Huh7, 6well | DV1-HNSB-P4(R1) | Infection of DV1-HNSB-P4 | 0.2 | 6.57 | 32.25 | 2.19 | 96hrs post infection | 10 | NA | **2.53** | **0.79** |
| | Huh7, 6well | DV1-HNSB-P4(R2) | Infection of DV1-HNSB-P4 | 0.2 | 4.81 | 23.59 | 1.60 | 96hrs post infection | 10 | NA | | |
| | Huh7, 6well | DV1-HNSB-P4-NS1(R1) | Transfection of DV1-HNSB-P4-NS1 | NA | 5.68 | NA | 1.89 | 96hrs post transfection | NA | 1 | **2.55** | **0.59** |
| | Huh7, 6well | DV1-HNSB-P4-NS1(R2) | Transfection of DV1-HNSB-P4-NS1 | NA | 6.65 | NA | 2.22 | 96hrs post transfection | NA | 1 | | |
| | Huh7, 6well | DV1-HNSB-P4-NS1(3.3) (R1) | Transfection of DV1-HNSB-P4-NS1 | NA | 12.48 | NA | 4.16 | 96hrs post transfection | NA | 3.3 | **4.36** | **0.33** |
| | Huh7, 6well | DV1-HNSB-P4-NS1(3.3) (R2) | Transfection of DV1-HNSB-P4-NS1 | NA | 12.54 | NA | 4.18 | 96hrs post transfection | NA | 3.3 | | |
| Fig2:(c) TUNEL | Vero, 6well | DV1-HNSB-P4(R1) | Infection of DV1-HNSB-P4 | 0.2 | 1.11 | 2.04 | 0.37 | 62 hrs post infection | 10 | NA | **0.40** | **0.09** |
| | Vero, 6well | DV1-HNSB-P4(R2) | Infection of DV1-HNSB-P4 | 0.2 | 0.98 | 2.04 | 0.33 | 62 hrs post infection | 10 | NA | | |
| | Vero, 6well | DV1-HNSB-P4(R3) | Infection of DV1-HNSB-P4 | 0.2 | 1.5 | 2.04 | 0.50 | 62 hrs post infection | 10 | NA | | |
| | Vero, 6well | DV1-HNSB-P4-NS1(R1) | Transfection of DV1-HNSB-P4-NS1 | NA | 2.7 | NA | 0.70 | 62hrs post Transfection | NA | 3.3 | **0.73** | **0.04** |
| | Vero, 6well | DV1-HNSB-P4-NS1(R2) | Transfection of DV1-HNSB-P4-NS1 | NA | 2.18 | NA | 0.73 | 62hrs post Transfection | NA | 3.3 | | |
| | Vero, 6well | DV1-HNSB-P4-NS1(R3) | Transfection of DV1-HNSB-P4-NS1 | NA | 2.32 | NA | 0.77 | 62hrs post Transfection | NA | 3.3 | | |

| Experiment with relevant Figure number | Type of cells & type of culture plate | Sample identification | Infection of DENV/ Transfection of NS1 | NS1 in Inocu-lum (µg) | Yield of NS1 (µg) excluding NS1 in inoculum | Fold change of NS1 (µg) over inoculum | Yield NS1 conc. (µg/ml) | Time of harvesting after infection or Transfection | Virus gE/cell | Transfected plasmid (µg) | Mean yield (µg/ml) | SD |
|---|---|---|---|---|---|---|---|---|---|---|---|---|
| Fig 2(e) TUNEL | Huh7, 6well | DV2-HNSB-P4(R1) | Infection of DV2-HNSB-P4 | 0.35 | 8.21 | 23.46 | 2.74 | 96hrs post infection | 10 | NA | **2.71** | **0.08** |
| | Huh7, 6well | DV2-HNSB-P4(R2) | Infection of DV2-HNSB-P4 | 0.35 | 8.33 | 23.80 | 2.78 | 96hrs post infection | 10 | NA | | |
| | Huh7, 6well | DV2-HNSB-P4(R3) | Infection of DV2-HNSB-P4 | 0.35 | 7.88 | 22.51 | 2.63 | 96hrs post infection | 10 | NA | | |
| | Huh7, 6well | DV2-HNSB-P4-NS1(3.3) (R1) | Transfection of DV2-HNSB-P4-NS1 | NA | 6.11 | NA | 2.04 | 96hrs post transfection | NA | 3.3 | **1.94** | **0.11** |
| | Huh7, 6well | DV2-HNSB-P4-NS1(3.3) (R2) | Transfection of DV2-HNSB-P4-NS1 | NA | 5.89 | NA | 1.96 | 96hrs post transfection | NA | 3.3 | | |
| | Huh7, 6well | DV2-HNSB-P4-NS1(3.3) (R3) | Transfection of DV2-HNSB-P4-NS1 | NA | 5.44 | NA | 1.81 | 96hrs post transfection | NA | 3.3 | | |
| | Huh7, 6well | DV3-HNSB-P4(R1) | Infection of DV3-HNSB-P4 | 0.23 | 3.85 | 16.74 | 1.28 | 96hrs post infection | 10 | NA | **1.21** | **0.15** |
| | Huh7, 6well | DV3-HNSB-P4(R2) | Infection of DV3-HNSB-P4 | 0.23 | 3.94 | 17.13 | 1.31 | 96hrs post infection | 10 | NA | | |
| | Huh7, 6well | DV3-HNSB-P4(R3) | Infection of DV3-HNSB-P4 | 0.23 | 3.13 | 13.61 | 1.04 | 96hrs post infection | 10 | NA | | |
| | Huh7, 6well | DV3-HNSB-P4-NS1(3.3) (R1) | Transfection of DV3-HNSB-P4-NS1 | NA | 5.2 | NA | 1.73 | 96hrs post transfection | NA | 3.3 | **1.56** | **0.16** |
| | Huh7, 6well | DV3-HNSB-P4-NS1(3.3) (R2) | Transfection of DV3-HNSB-P4-NS1 | NA | 4.58 | NA | 1.53 | 96hrs post transfection | NA | 3.3 | | |
| | Huh7, 6well | DV3-HNSB-P4-NS1(3.3) (R3) | Transfection of DV3-HNSB-P4-NS1 | NA | 4.23 | NA | 1.41 | 96hrs post transfection | NA | 3.3 | | |
| Fig:3(a) Ladder assay | Vero, T-25 | DV1-HNSB-P4 | Infection of DV1-HNSB-P4 | 0.49 | 2.64 | 5.39 | 0.53 | 62 hrs post infection | 10 | NA | | |
| | Vero, T-25 | DV1-HNSB-P4+Camp | Infection of DV1-HNSB-P4 | 0.49 | 2.72 | 5.55 | 0.54 | 62 hrs post infection | 10 | NA | | |
| Fig:3(c) DAPI | Huh7, 6well | DV1-HNSB-P4(R1) | Infection of DV1-HNSB-P4 | 0.32 | 3.05 | 9.53 | 1.02 | 48hrs post infection | 10 | NA | **0.95** | **0.08** |
| | Huh7, 6well | DV1-HNSB-P4(R2) | Infection of DV1-HNSB-P4 | 0.32 | 2.6 | 8.13 | 0.87 | 48hrs post infection | 10 | NA | | |
| | Huh7, 6well | DV1-HNSB-P4(R3) | Infection of DV1-HNSB-P4 | 0.32 | 2.89 | 9.03 | 0.96 | 48hrs post infection | 10 | NA | | |
| | Huh7, 6well | DV1-HNSB-NS1(R1) | Transfection of DV1-HNSB-NS1 | NA | 3.51 | NA | 1.17 | 48hrs post transfection | NA | 1 | **1.15** | **0.03** |
| | Huh7, 6well | DV1-HNSB-NS1(R2) | Transfection of DV1-HNSB-NS1 | NA | 3.5 | NA | 1.17 | 48hrs post transfection | NA | 1 | | |
| | Huh7, 6well | DV1-HNSB-NS1(R3) | Transfection of DV1-HNSB-NS1 | NA | 3.34 | NA | 1.11 | 48hrs post transfection | NA | 1 | | |

**Table S1. Estimation of NS1 yields and concentrations from different experiments**

Secreted NS1 levels in experiments were measured using semi-quantitative NS1 ELISA as described in methods. Individual ELISA, with standards, was performed for each experiment. In case of infection with DV1-HNSB-P4, DV2-HNSB-P4 or DV3-HNSB-P4, increase in sNS1 yield was expressed in micrograms and as fold-increase over sNS1 level present in the inoculum. Different types of experiment are indicated next to the respective figure numbers.
